# The thalamic reticular nucleus provides spatially and temporally tuned inhibition to the lateral geniculate nucleus across species

**DOI:** 10.64898/2026.11.10.763754

**Authors:** Yizhan Miao, Ulas M. Ciftcioglu, Alexis S. Gorin, Rahi Dakwala, Seohee Ahn, Friedrich T. Sommer, Judith A. Hirsch

## Abstract

By proposing a role in spatial attention, Francis Crick piqued interest in the thalamic reticular nucleus (TRN), a GABAergic structure that receives input from relay cells projecting to cortex and feeds back inhibition in return. Importantly, predictions of Crick’s hypothesis extend beyond attention to suggest that TRN provides selective, localized inhibition to relay cells and, thus, influences sensory processing *per se*. To ask if this latter role for TRN holds across species, we compared previous recordings from cat with a novel dataset from mouse, focusing on vision. We estimated receptive field size and structure of reticular cells in the visual sector of TRN and of relay cells in the dorsal lateral geniculate nucleus (dLGN). Cells in both nuclei encoded specific visual features and had receptive fields of comparable size within binocular visual space. Moreover, paired recordings from both structures suggested that TRN is able to promote burst firing in dLGN in addition to modulating spike rate. Thus, TRN is not only important for attentional regulation, but is also able to make fine-tuned contributions to early sensory processing.

## Introduction

En route to cortex, thalamic axons contact the thalamic reticular nucleus (TRN) which inhibits relay cells in return^1,2^. Here we ask how this feedback inhibition might impact sensory processing and approach the question by studying the region of TRN that lies within the early visual pathway and connects with the dorsal lateral geniculate nucleus of the thalamus (dLGN). Our first goal was to understand the qualities of inhibition that TRN provides. Next, we considered how this inhibition engages with relay cells to alter their activity patterns. Towards these goals, we recorded visual responses of reticular and relay cells across species, using an existing dataset from cat and obtaining new recordings from mouse.

For highly visual mammals like cat and monkey, the representation of the visual field is mainly binocular because their eyes face forward. The receptive fields of most retinal ganglion cells in these species have a similar (center-surround) shape, are smallest within a specialized region that represents the center of gaze (e.g. the fovea), and grow larger with eccentricity^3^. Rodents are different. Their eyes point outwards such that the binocular component of the visual field is limited to a small central frontal area^4^. As well, there are many different types of retinal ganglion cells, with commensurately varied receptive fields and response properties, that are distributed in a nonuniform fashion that reflects the animal’s ecological niche^5–11^. For example, even though rodent lacks a strict foveal homolog and only a small subset of ganglion cells have center-surround receptive fields, these are concentrated in the retinal region that encodes binocular space^12^ and is crucial for visually demanding tasks like prey capture^13,14^. This finding suggests conserved principles of thalamic visual processing across species.

Receptive fields in mammalian dLGN largely mirror those in retina^15–18^ and relay cells provide driving input to TRN^19,20^. In cat, TRN cells have complex visual features quite different from the center-surround structures preferred by relay cells^21^. Moreover, receptive field size in both cat TRN and dLGN is comparable at any given eccentricity in the visual field^22^. While receptive field eccentricity (i.e., spatial position with respect to a single central coordinate) in highly visual animals maps nicely onto the functional organization of retina (e.g. foveal versus peripheral), this is not the case for rodent. Thus, we assigned results from mouse to a binocular/monocular visuotopic reference frame.

Within the binocular zone, the average receptive field size in TRN was comparable to that for dLGN, though larger in the monocular zone. Further, murine receptive field types in TRN were at least as diverse as those in dLGN. These results suggest that spatially precise and feature selective feedback inhibition from TRN is important for form vision across species.

To explore how this feedback inhibition influences response dynamics in dLGN, we considered that neurons in both nuclei fire in two modes, tonic trains of spikes or rapid bursts driven by calcium channels that require substantial hyperpolarization to reset^23,24^. Bursting is normally associated with sleep but stimuli that drive strong inhibition can evoke bursts as well, as demonstrated in dLGN^25–28^. We found that this was also the case in TRN for both species and show how reticular bursts might contribute to priming bursts in relay cells (local interneurons also contribute to this process^27^).

Drawn in by its unique position between thalamus and cortex and propensity towards burst firing, Francis Crick proposed a role for TRN in spatial attention^29^; studies in awake behaving animals support this view^30,31^. Our finding that TRN is able to supply feature selective, localized inhibition to relay cells across species provides mechanistic support for Crick’s idea. Further, synaptic connections within murine TRN link cells that project to first-order thalamic nuclei with those that project to higher-order nuclei^32,33^, and there is evidence that this could be true in cat and primate as well^34,35^. Our observation that visual stimuli drive bursts in TRN and that these, in turn, might promote bursting in relay cells is particularly interesting in the context of the recent discovery that bursts in pulvinar drive spatial attention^36^. Thus, TRN is poised to contribute to diverse aspects of sensory function.

## Results

To understand how inhibition from the visual sector of TRN might influence information that relay cells in dLGN send to cortex, we recorded visual responses from relay cells in dLGN (n=617) and neurons in TRN (n=184) in 118 adult mice (Figure 1); some relay cells were included in earlier studies^17,37^. To compare responses in murine thalamus with those in carnivore, we reanalyzed responses from relay cells (n=61) and TRN cells (n=44) recorded from adult cat^21,22^.

**Figure 1:**
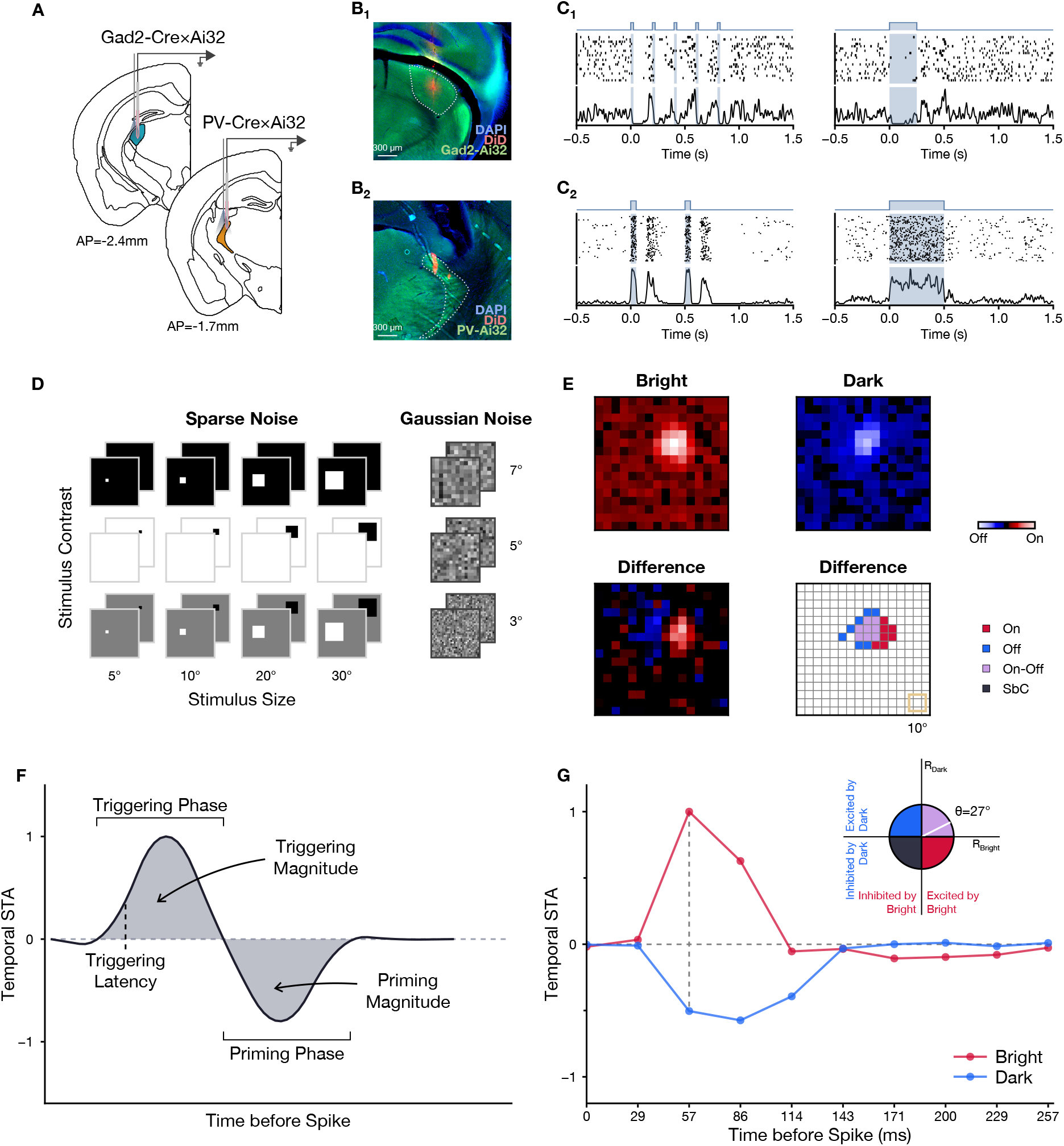
Estimating receptive fields of relay cells in dLGN and cells in TRN using *in vivo* recordings. (**A**) Extracellular recordings targeting dLGN and/or TRN were made in anesthetized transgenic mice (Gad2-Ai32, dLGN; PV-Ai32, TRN); relay cells and reticular cells were identified optogenetically. **(B**) Example micrographs show electrode tracks (DiD, red) through dLGN (B_1_) and through TRN (B_2_). **(C**) Rasters show how pulses of blue light (blue trace and shading) evoked distinct responses from a relay cell in dLGN (C_1_) and a cell in TRN (C_2_). C_1_: Pulses of blue light (left: 25ms at 5Hz, right: 100ms at 1Hz) led to periods of suppression followed by rebound bursts. C_2_: Pulses of blue light (left: 50 ms at 2 Hz, right: 250 ms at 1 Hz) drove spikes in TRN cells, often followed by cycles of suppression and firing. **(D**) Visual stimuli. Left: sparse noise (black and white squares of varying size and contrast shown randomly on a 16×16 grid, sized 80°×80°). Right: Gaussian noise of different pixel sizes. **(E**) Peak frames of STAs from an On-Off relay cell mapped with sparse noise. Top row: On and Off responses were estimated using only bright (red) or dark (blue) stimuli, respectively. Bottom left: difference map made by subtracting the dark from the bright STA; On-Off responses in mouse are often asymmetrically strong such that only the dominant contrast for each pixel is reflected in difference map; overall bright pixel values dominated dark ones in the case of this frame. Bottom right: A spatial “footprint” showing the spatial extent of both On responses and Off responses and the region of overlap; yellow square indicates stimulus size (10°). **(F**) Schematic of a stereotypic biphasic temporal STA (tSTA): a “triggering phase” that evokes a spike is preceded by a “priming phase” that prepares the membrane to fire. “Triggering latency” is defined as earliest time at which response amplitude reaches a statistically significant level. “Triggering magnitude” and “priming magnitude” are the area of each phase. **(G**) tSTAs of the On-Off relay cell from E (see <u>METHODS</u>); triggering latencies of both bright and dark tSTAs were 57ms (dashed vertical line). Inset: Bright-dark polarity score (white, *θ* = 27°) shown along a unit circle.

### Cells in murine TRN have receptive-field types as diverse as those of relay cells in dLGN

In cat, TRN provides novel forms of spatially discrete inhibition to dLGN^21,22^. We asked if there were an analogous role for murine TRN, where relay cells themselves have diverse types of receptive fields (Figure 2A). The most single common type of receptive field in murine dLGN is center-surround (i.e., “On-center Off-surround” (On-Center) or “Off-center On-surround” (Off-Center)), with the remaining population comprising On, Off, On-Off, and Suppressed-by-Contrast (SbC) profiles (Figure 2B)^16,17,37,38^.

**Figure 2:**
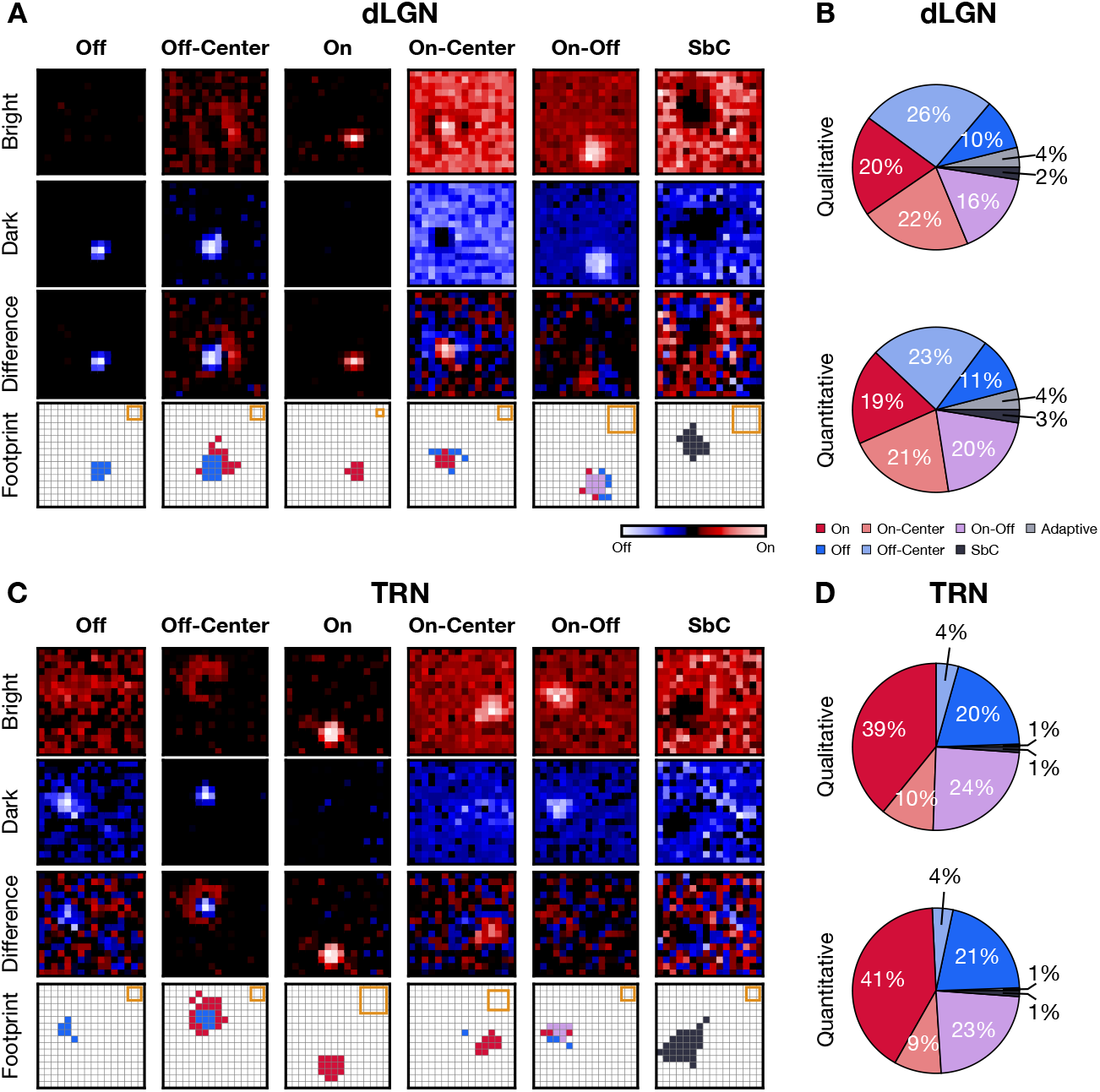
Diverse receptive-field types in murine dLGN and TRN. (**A**) Peak frames of bright, dark, and difference STAs above corresponding spatial footprints from Off, Off-Center, On, On-Center, On-Off, and Suppressed-by-Contrast (SbC) cells in dLGN; Yellow squares indicate stimulus size; grid (gray) spacing, 5°. **(B**) Pie-charts show distributions of different receptive-field types of cells in dLGN categorized qualitatively (top, n=617) or quantitatively (bottom, n=589) by the bright-dark polarity score (see <u>METHODS</u>). **(C**) Example responses of all types of receptive fields in TRN, conventions as in (A). **(D**) Distributions of different types of receptive fields in TRN, conventions as in (B); top, qualitative categories (n=184), bottom, quantitative categories (n=175).

Because receptive fields in TRN can be patchy and irregular, we developed a non-parametric “footprint” method^22^ to estimate the size of the spatiotemporal receptive field in cat and adapted this method for the current study. Using standard qualitative criteria^16,17^ to assign receptive field types from the footprints, we found that (Figure 2C) On, Off, and On-Off cells were most common; surprisingly, there was also a small population of center-surround cells (Figure 2C, D). We then used a quantitative method called the bright-dark polarity score^17,37^ to sort receptive field types into four main categories: On, Off, On-Off and SbC. The first step in this process was to use the footprint as a spatial mask to calculate the time course of the STA (tSTA, see METHODS; Figure 1F) from the bright and the dark maps obtained using sparse noise. The second step was to calculate the bright-dark polarity score from the response at the start of the triggering phase (the stimulus that directly evokes a spike) (Figure 1G). Surrounds, if present, were assessed by eye. Both quantitative and qualitative results were similar and show that receptive fields in mouse TRN were as diverse as in mouse dLGN, although, at least for our sample size, the distribution was different (Figure 2B, D).

### Receptive fields in murine visual thalamus scale with position in visuotopic space

In highly visual animals, receptive-field sizes are smallest at the center of gaze (near the area centralis in carnivore or the fovea in primate) and scale with eccentricity. Indeed receptive fields in dLGN and TRN are of comparable size at a given eccentricity at all^39^ but the most peripheral region of visual space, where reticular fields grow disproportionately large^22^. Although mouse does not have a fovea or area centralis *per se*, the distribution of different types of ganglion cells varies according to retinal position^5,6,40^ and there is accumulating evidence that receptive field sizes are smaller in the binocular zone (near the center of the frontal visual field)^12^. Therefore, we explored differences in receptive field type and size in dLGN and TRN across visual space. The visuotopic location of the receptive field was determined by projecting the footprint onto the left visual field, with the receptive field center defined by the centroid of the footprint. (Figure 3A, also see METHODS). Although, technically, it is not possible to sample each structure evenly, our dataset covered a wide range of visuotopic space, including recordings as far lateral as 120° along the azimuth: dLGN (Figure 3B) and TRN (Figure 3C).

**Figure 3:**
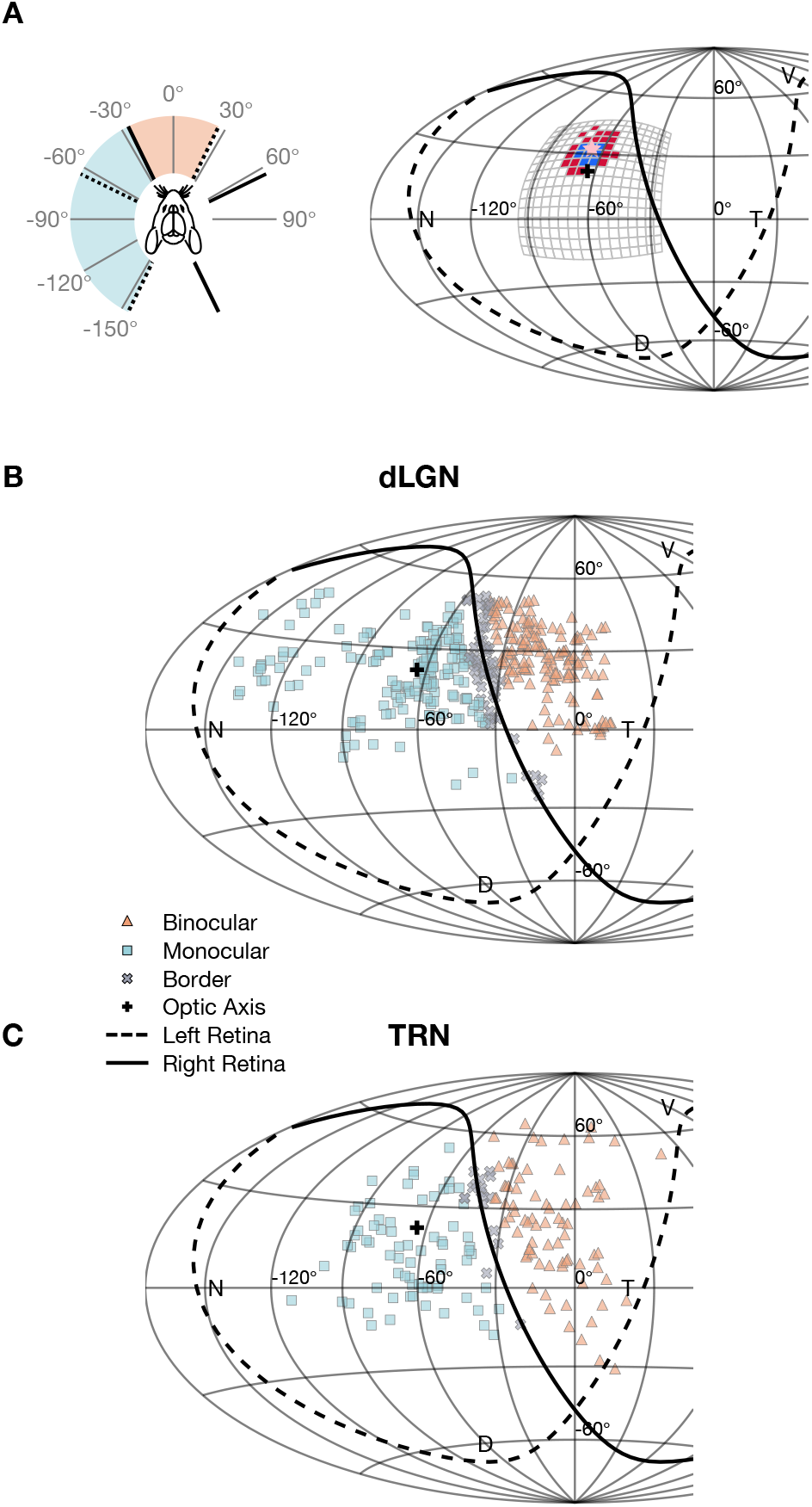
Sampling receptive fields from visual thalamus across visuotopic space. (**A**) Schematic representation of receptive field placement within visuotopic space. Left: The left visual field is divided into binocular (peach), monocular (blue), and border regions (5° around the right visual field limits, gray). The extent and the optic axis of the left visual field are delimited with dashed lines and that of the right visual field with solid lines; optic axis: ±64° azimuth and 22° elevation^4^. Right: A representative footprint of an Off-Center relay cell projected on visuotopic space (Hammer projection), with the receptive-field center determined by the footprint centroid, pink star. Dashed and solid lines show the border of the left and right visual fields, respectively; N, nasal; T, temporal; D, dorsal; V, ventral. Spacing between each grid line, 30°. **(B,C**) Visuotopic distributions of receptive-field centers of sampled dLGN cells (B, n=378) and TRN cells (C, n=167).

At first glance, our results suggest a general trend for footprint sizes to be smallest in the binocular zone and increase with moving deeper into the monocular visual field for both dLGN (Figure 4A) and TRN (Figure 4B). To quantify this change, we calculated the angular distance (see METHODS) from each footprint center to the border. Consistent with our initial impression, there was significant linear relationship between footprint size and angular distance (Figure 4C, D).

**Figure 4:**
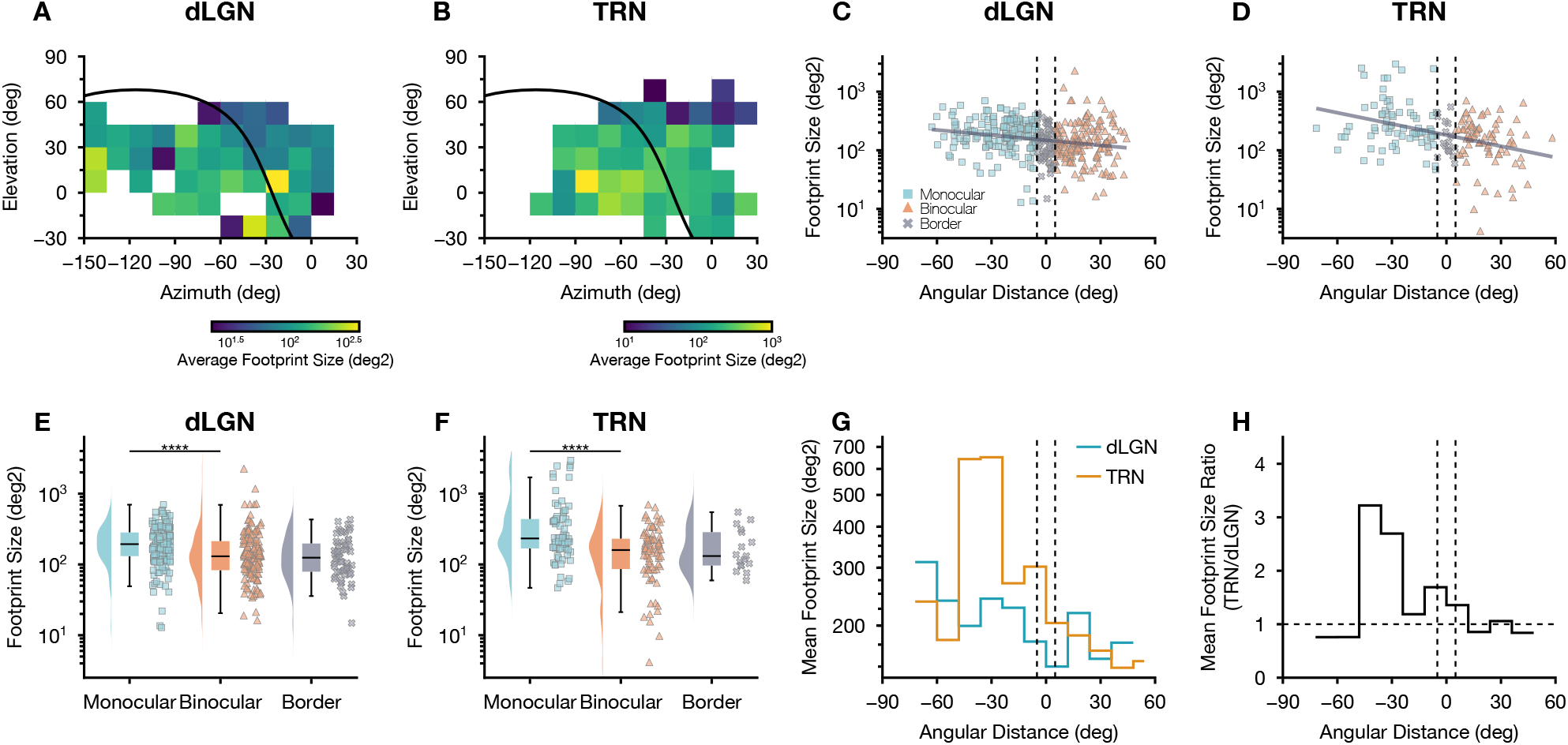
Non-uniform distribution of receptive-field sizes in murine dLGN and TRN across visuotopic space. (**A, B**) Heatmap of averaged footprint sizes (all significant pixels; see <u>Figure S2</u> for dominant-pixel sizes) across visuotopic space (bin size, 15°) for cells in dLGN (A) and TRN (B) from Figure 3. Solid line, border of right retina (i.e., border of binocular/monocular visual space). **(C, D**) Scatter plots show the relationship of footprint sizes from dLGN (C) and TRN (D) plotted on a log scale against the angular distance from the border of the monocular visual field, with the linear regression for each population plotted as solid line. Vertical dashed lines indicate the border between binocular and monocular zones. (F-test; dLGN, p=1.810 × 10^−5^; TRN, p=1.977 × 10^−7^). **(E, F**) Raincloud plots show ranges of footprint sizes in the monocular, binocular, and border regions for cells in dLGN (E, Mann-Whitney U test, p=7.596 × 10^−7^) and TRN (F, Mann-Whitney U test, p=6.120 × 10^−5^). **(G**) Histograms plot distributions of averaged footprint sizes of cells in dLGN and TRN along the angular distance from the border of binocular/monocular visual field; bin size, 12°. **(H**) Histogram of ratios between the averaged footprint size of dLGN cells and TRN cells at each angular distance. The horizontal dotted line indicates ratio=1.

Since the footprint analysis excludes components of the receptive field that do not reach statistical significance, regions that respond only weakly to small stimuli, such as the surround, are not always captured. Therefore, we repeated our comparison of footprint size to visuotopic position using only regions of the receptive field driven by the preferred, or dominant, stimulus contrast (i.e., dark or bright) (see METHODS). The results were the same as those obtained using the full footprint (Figure S2).

The binocular component of the visual field recalls areas of greatest acuity in highly visual animals. Thus, we compared footprint sizes in the binocular zone (angular distance > 0°) to those in the monocular zone (angular distance < 0°). To minimize the effect of any potential measurement errors, we excluded the samples that fell within ± 5° of the border between regions. The mean footprint sizes within the binocular zone were significantly smaller than those of the monocular region in both dLGN (Figure 4E, Mann-Whitney U test, p=7.596 × 10^−7^; binocular, n=150; monocular, n=170) and TRN (Figure 4F, Mann-Whitney U test, p=6.120 × 10^−5^; binocular, n=78; monocular, n=72).

Next, we asked if receptive-field sizes of cells in dLGN and TRN were comparable, as they are in cat^22^ and seem to be in macaque^39^. Our results show that averaged footprint sizes in TRN and dLGN were similar in the binocular zone, but that the mean footprint size in TRN far exceeds that for dLGN in the monocular zone (Figure 4G). We generated a histogram for the ratios of footprint sizes in TRN and dLGN, which shows that values were, indeed, close to one within the binocular zone and grew progressively larger within the monocular field (Figure 4H). Thus, our results show that receptive fields in murine dLGN and TRN are of comparable size and are smaller in the slice of visual space that both eyes see than in the monocular visual field.

### Visual responses: burst and tonic modes

Cells in TRN and dLGN fire in two modes—burst and tonic. Bursts are mediated by a family of low-threshold activated T-type calcium channels that require exceptionally strong hyperpolarization to reopen^23,24^. As we have illustrated earlier, the time course of the receptive field (tSTA) is often biphasic, comprising a “priming” phase that, presumably, hyperpolarizes the membrane in preparation for firing, and a “triggering” phase that directly evokes a spike. The visual features encoded by tonic or burst spikes usually have similar spatial extents; however the priming phase of the response is more prominent in burst than tonic firing in dLGN^27,28,41^. Here we asked if there were parallel properties in TRN by comparing tSTAs that evoke tonic versus burst spikes in dLGN and TRN from cat (cat dLGN, n=59; cat TRN, n=41) and mouse (mouse dLGN, n=164; mouse TRN, n=38), focusing on cells with biphasic responses.

The spatial receptive fields associated with both firing modes are depicted in the frame-by-frame evolution of the STA for both dLGN and TRN (Figure 5); only the cardinal spike of each burst was used for the analysis^21^. As in dLGN, the spatial extents of burst and tonic spikes in TRN were similar. To further quantify the magnitude of the two phases of the tSTA, we used the normalized response within the extent of the footprint. Again, the priming phase appears to be proportionally stronger for bursts than for tonic spikes (Figure 5)—not only for dLGN but for TRN as well.

**Figure 5:**
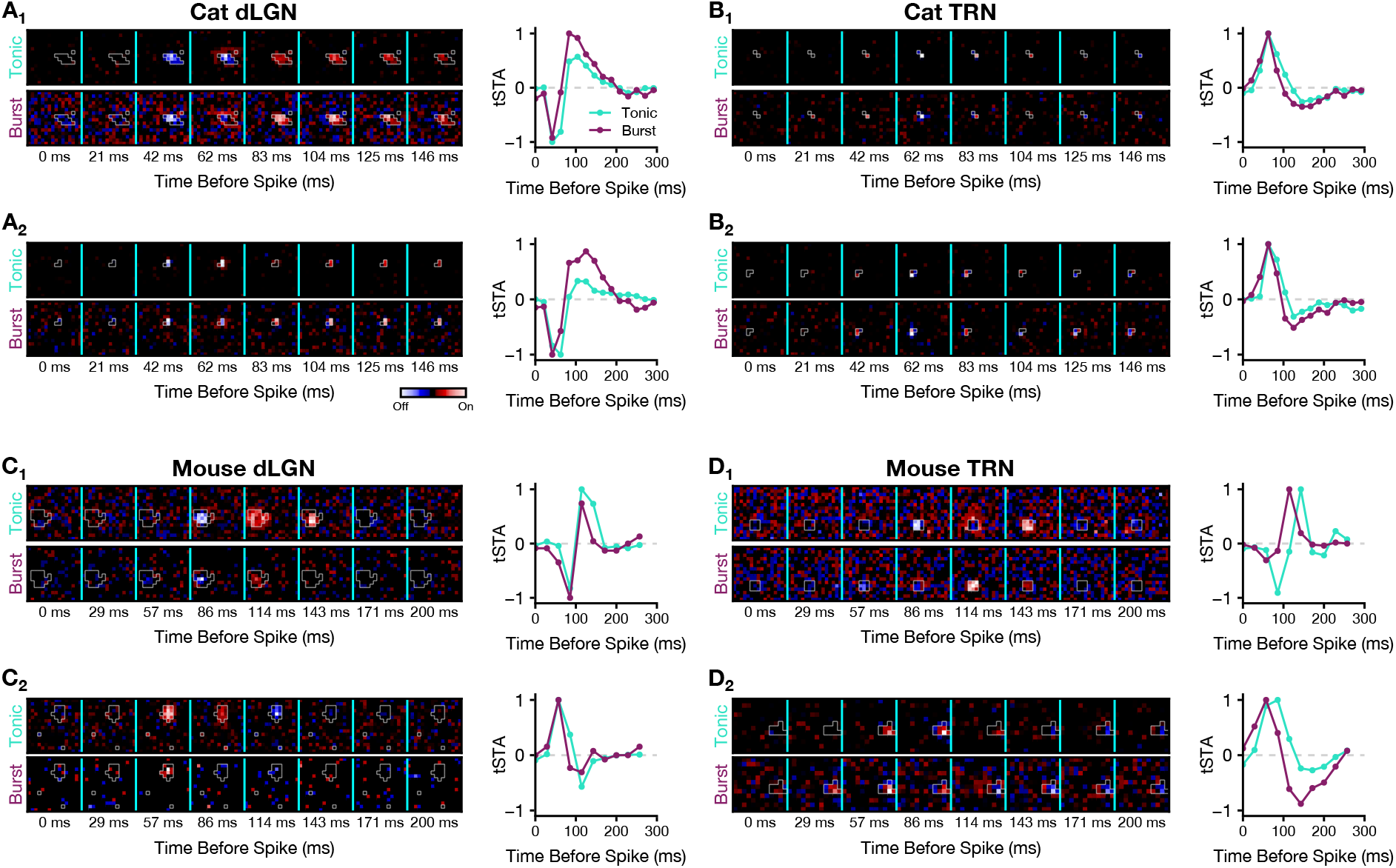
Estimating spatiotemporal receptive fields from tonic and burst spikes. (**A**) Receptive-field estimations for two example cat relay cells in dLGN. Left: The spatial and temporal evolution of the STAs for tonic spikes (top) and the cardinal spike of each burst (bottom). Gray contours outline the corresponding footprint for each cell. Right: Normalized tSTAs computed by summing pixel values within the footprint show that the priming phase before bursts is proportionally larger than that for tonic spikes; positive values, response to bright stimuli (On), negative values, response to dark stimuli (Off). **(B, C, D**) Conventions as in (A) but for cells in cat TRN (B), cells in mouse dLGN (C) and TRN (D).

To quantify the differences in tSTA profiles for burst and tonic spikes, we defined a “priming-triggering ratio” (see METHODS), which assesses the relative magnitude (area under the curve) of the priming versus the triggering phase. The priming-triggering ratios were higher for bursts than those for tonic spikes, especially for dLGN of both species, cat (Figure 6A, Wilcoxon signed-rank test, p=2.382 × 10^−9^) and mouse (Figure 6B, Wilcoxon signed-rank test, p=2.295 × 10^−8^). The result in cat TRN was similar (Wilcoxon signed-rank test, p=0.014), and somewhat more robust than that for mouse (Wilcoxon signed-rank test, p=0.034). Together, these results suggest that, across species, burst and tonic spikes both in dLGN and TRN encode visual features with similar spatial extents but distinct temporal profiles.

**Figure 6:**
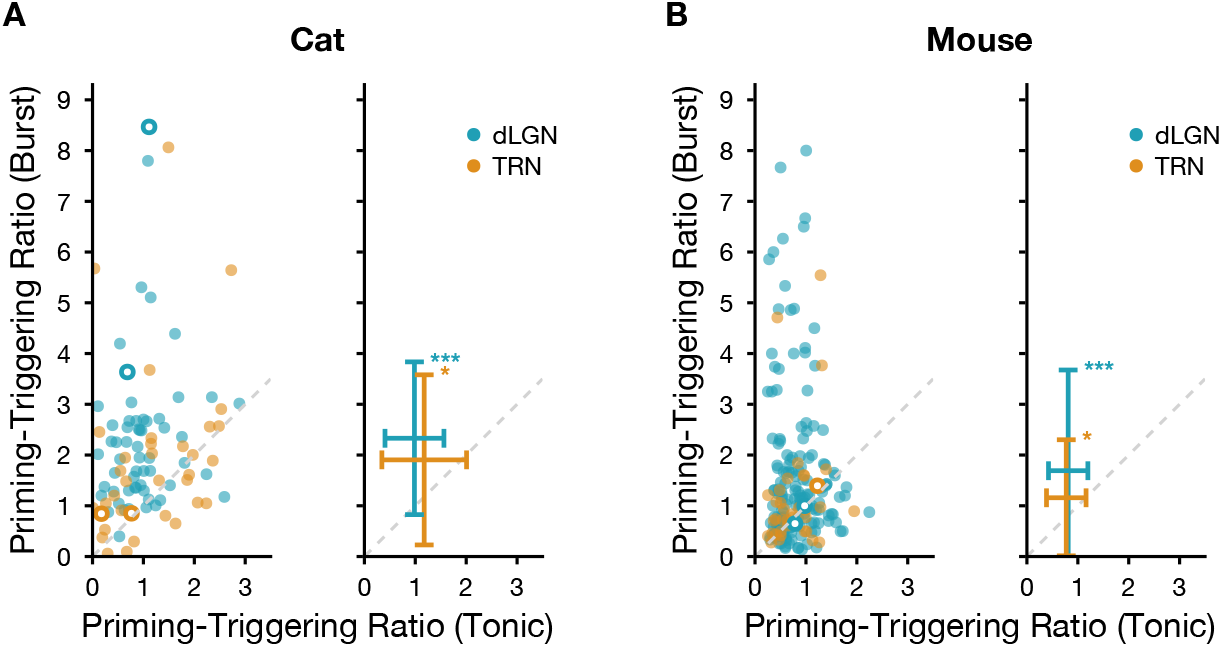
Priming-triggering ratio between tonic and burst spikes from dLGN and TRN. (**A**) Left: Scatter plots of the priming-triggering ratio (see <u>METHODS</u>) for tonic versus burst spikes for cells in cat dLGN (n=59) and TRN (n=41); dashed diagonal, unity line. Right: Plots of the mean and SD ratio for each group. Open circles highlight the examples shown in Figure 5. **(B**) Conventions as in (A), but for cells in mouse dLGN (n=164) and TRN (n=38). Wilcoxon signed-rank test, cat dLGN p=2.382 × 10^−9^, cat TRN p=0.014, mouse dLGN p=2.295 × 10^−8^, mouse TRN p=0.034.

### Bursts from TRN might help prime bursts in dLGN

We were interested in exploring how bursts in TRN might influence visual processing in dLGN. Our past work in cat provided evidence that inhibition from local interneurons can help prime bursts in dLGN^27^. We asked whether bursts in TRN might promote bursting as well. Thus, we made simultaneous recordings in mouse dLGN and TRN (Figure 7) and were able to monitor several pairs of cells that encoded overlapping regions of visual space. We presented natural movies to these cells as the statistics of these stimuli are well suited for driving bursts (Figure 7A)^26,27,42^, and also help illustrate patterns of neuronal activity as animals navigate their environment. Then, we extracted trains of tonic spikes or trains made only from the cardinal spike of each burst, as above^21^, and cross-correlated results for each pair of cells. The cross-correlogram for an example On-Off cell in dLGN and an On-Off cell in TRN (Figure 7B) shows a peak near zero for tonic spikes. The cross-correlogram of cardinal spikes in bursts, however, showed a systematic shift towards negative lags (peaks are at least 2σ above baseline), suggesting that bursts in TRN precede those in dLGN. The same overall pattern of response was also seen for a second example pair—an On-Off dLGN and Off TRN cell (Figure 7C), with TRN bursts leading dLGN bursts. Cells at displaced positions in visual space lacked such correlations (Figure S3B). Of course, these results do not consider how milder forms of inhibition from TRN also influence firing and we hope to investigate this in the future.

**Figure 7:**
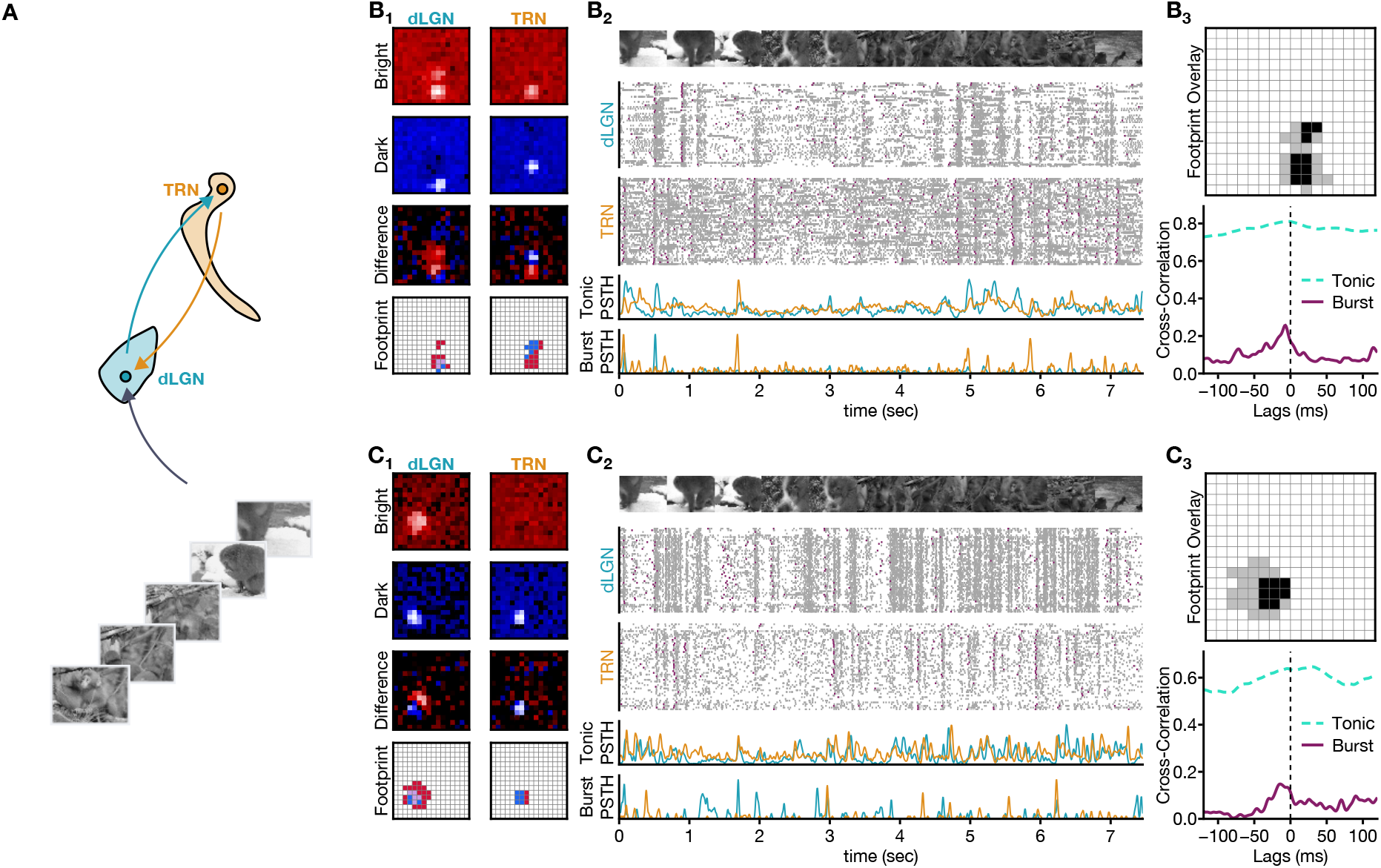
Bursts in TRN cells preceded bursts in relay cells. (**A**) Schematic illustrating simultaneous extracellular recordings from dLGN and TRN made during the presentation of naturalistic movies. **(B**) Example responses from a pair consisting of a dLGN On-Off cell and a TRN On-Off cell: **(B_1_**) Peak frames of STAs and corresponding spatial footprints of the example pair of cells. **(B_2_**) Rasters show responses to multiple trials of the naturalistic movie (shown below representative frames of the stimulus). The cardinal spike of each burst is shaded pink. Bottom: PSTHs of tonic spikes and cardinal spikes. **(B_3_**) Top: union footprint shows that the receptive fields of the two cells are largely overlapped (black, overlapped pixels). Bottom: cross-correlations of tonic and burst PSTHs; dLGN responses were used as reference. Tonic cross-correlation peaked around zero with no lag; burst cross-correlation peaked with a negative time lag. Vertical dashed line, lag = 0ms. **(C_1_-C_3_**) Conventions as in B, but for a different pair; a dLGN On-Off cell and a TRN Off cell.

## Discussion

Our results revealed that basic attributes of TRN that had been observed in highly visual animals were conserved in mouse. For example, receptive fields in murine TRN were selective for diverse visual features and were similar in size to those in dLGN within the binocular zone. Thus, across species, TRN is able to provide tailored suppression to dLGN in the region of visual space where the animal actively directs its gaze, complementing local inhibition from interneurons. Notably, reticular cells differ from local interneurons in that they fire in both tonic and burst modes^24^, much like relay cells. We found that visual stimuli drove bursts in TRN, as has been long documented in dLGN^26,28^. Further, cross-correlation analysis of simultaneous recordings from retinotopically aligned locations in TRN and dLGN suggested that reticular input could facilitate bursting in relay cells. These results might also have bearing on activity in higher-order nuclei. TRN cells that project to first-order thalamic nuclei, such as dLGN, share electrical^32^ and chemical^33^ synapses with neighbors coupled with higher-order nuclei such as pulvinar.

### Connections of TRN with first-and higher-order thalamic nuclei

TRN comprises adjacent subnetworks that connect with first or higher-order thalamus. Inputs to first-order TRN are tightly topographic while those innervating higher-order TRN are more diffusely distributed (primate^34,43^; carnivore^44^; lagomorph^45,46^; rodent^47–50^). Further, reticular cells within first-order subnetworks produce more bursts than their higher-order counterparts^47,51^. In mouse, the pulvinar-coupled region of TRN forms a very thin “edge” that wraps around a thicker “center” that is linked to dLGN^47,48,51,52^. Thus, it is likely that we sampled at least a few edge cells as the electrode passed through TRN. As yet, however, we have not found evidence for a significant bimodal distribution of responses based on either receptive field size or burst statistics and so aggregated all our recordings from murine TRN prior to analysis.

### Receptive field organization in visual thalamus

Highly visual animals rely on binocular vision, with the sharpest acuity achieved at specialized retinal regions that encode the central visual field—the fovea in primate and the area centralis in cat, for example. This organization is preserved along the early visual hierarchy, such that receptive-field sizes are smallest centrally and scale with eccentricity^22,53,54^. Although mouse does not have a fovea *per se*, ganglion cell types are nonuniformly distributed across retina^5,6^. For instance, there is a region of the central binocular visual field that has a concentration of cells with relatively small^12^ center-surround receptive fields^7,8^. These are among the cohort of ganglion cells that project to dLGN and are important for existential behaviors such as prey pursuit and capture^13,14,55^.

We thus separately analyzed receptive fields within the binocular zone from those that lie in the monocular visual field to evaluate potential regional differences. To compute receptive field size, we used an unbiased, non-parametric method^22^. In dLGN, average receptive field size in the binocular zone was significantly smaller than in the monocular zone. There was also a trend for receptive fields to grow larger from the center of visual field towards the periphery. A previous study of dLGN that focused on the distribution of color preference found no difference in receptive field size as a function of retinotopic position, but that dataset was mainly obtained from the binocular zone^56^. Lastly, the distribution of receptive field size we observed in dLGN is likely relayed to cortex, where population imaging^57^ in mouse V1 reveals a trend towards smaller receptive field sizes in the binocular visual field.

Companion recordings from TRN revealed that receptive field sizes in the binocular field are similar in size to those in dLGN and significantly smaller than those in monocular regions. Further, the scale of reticular receptive fields increased disproportionately in the monocular versus the binocular zone, such that the ratio of receptive field sizes in TRN versus dLGN nearly doubled. An earlier study found that average receptive field size in TRN was substantially larger than that in dLGN^58^. These data, however, were aggregated across TRN and included anatomical regions representing the far periphery; our findings likely explain the discrepancy. Thus, evaluation of receptive field size across space emphasizes regional specializations.

Assessing differences in receptive field complexity between dLGN and TRN proved more challenging for mouse than for cat. In cat, reticular cells are selective for complex visual features^21^, very different from the stereotyped center-surround profiles common across relay cells and interneurons in dLGN^59,60^. By contrast, there is a great deal of receptive field diversity in mouse dLGN^16,17^. That said, our results revealed that neurons in TRN had receptive fields at least as diverse as those in dLGN, including the surprising presence of center-surround shapes. It is unlikely that we confused TRN for either division of LGN as we used a combination of optogenetic identification, histological confirmation of the electrode track, and spike statistics to determine recording sites.

As yet, our sample size remains too small to make definitive statements about receptive field distribution across visual space. Also, we mapped only the contralateral receptive fields in our study, so are unable to report how many receptive fields received binocular input. That said, there is evidence to suggest that single reticular cells could have a binocular influence, as some project to multiple layers in dLGN^43,61,62^. All told, our impression from the literature and our own results suggest that TRN provides feature-selective inhibition to support behaviors requiring higher visual acuity or focused spatial attention across species.

### Role of bursts in reticulogeniculate connections

Much like relay cells, reticular cells fire T-type channel initiated bursts of spikes in addition to tonic trains of spikes, though the kinetics of the channels in TRN produce lower threshold, longer, and slower bursts^24,63,64^. To explore the possibility that bursts in TRN might be visually driven, we used a toolkit based on studies of dLGN^21,28^. We confined our analysis to cells with biphasic spatiotemporal receptive fields—in which an initial triggering phase that elicits firing is preceded by a priming phase that prepares the membrane to spike. In TRN, as had been found in dLGN^28^, bursts were evoked by stimuli that had a similar shape but a stronger priming phase than stimuli that preceded tonic firing. Further, cross-correlation of visual responses made from simultaneously recorded relay and reticular cells with overlapping receptive fields revealed that bursts in TRN often began before those in dLGN. Hence, inhibition from TRN has the potential to facilitate bursts in relay cells and thereby impact signals sent to cortex^58^.

Even though the region of TRN we sampled projects mainly to dLGN, reticular cells interact across subnetworks, with overlap between first– and higher-order TRN evident in primates including marmoset^34^, galago^62^ and macaque^34^, cat^35,65^ and mouse^32,33,66^. The mechanism of this interaction has been explored best in mouse, where cells in first– and higher-order subnetworks cross the divide to synapse with each other^32,33,66,67^. Thus, through such network interactions, primary visual TRN might contribute to bursts in pulvinar, which recent work in monkey shows are involved in attention^36^. More broadly, because first– and higher-order divisions of TRN are involved in different cortical feedback loops^68–70^, interactions among reticular cells in different subnetworks stand to have diverse impacts on thalamocortical dynamics^71^.

## Materials and Methods

### Experimental Models and Subject Details

#### Animals

All procedures were performed in accordance with the U.S. National Institutes of Health and approved by the Institutional Animal Care and Use Committees of the University of Southern California. All subjects were mice aged ≥ 8 weeks to avoid the visual critical period. Subjects were the progeny of either Gad2-IRES-Cre mice (Jackson Lab, RRID:IMSR_JAX:028867) or Pvalb-IRES-Cre mice (Jackson Lab, RRID:IMSR_JAX:017320) crossed with Ai32 (ChR2(H134R)/EYFP, Jackson Lab, RRID:IMSR_JAX:024109) mice (i.e., Gad2-Ai32 and PV-Ai32). There was no detectable difference between results obtained with males and females, so data from both sexes were pooled.

### Experimental Design

#### Electrophysiological recordings

Prior to electrophysiological recording, animals were given an injection of chlorprothixene (1 mg/kg, i.p.) after which anesthesia was initiated and maintained with urethane (0.5 g/kg – 1 g/kg, i.p.) or with isoflurane (0.5%). The skull was exposed and affixed to a headpost. A craniotomy was made over dLGN or TRN, with a grounding head screw secured over the contralateral hemisphere or cerebellum. Wound margins were infiltrated with lidocaine (0.5%) and body temperature was maintained at 37 °C.

Extracellular recordings were made with multi-conductor multi-shank electrodes (Cambridge NeuroTech) that were connected to a digital multiplexing headstage and a Digital Lynx 4SX-M acquisition system (Neuralynx Inc.); high– and low-pass digital filters were set to 0.5 Hz and 7 kHz and the sampling rate was 30 kHz. Electrode tracks were registered histologically using a red-shifted lipophilic dye (DiD, Invitrogen catalog#V22887) applied to the electrodes before insertion or, when the dye was undetectable, from the path created by the electrode.

Recordings from earlier studies were sometime obtained using glass pipettes in whole-cell or cell-attached mode, as described previously^17,37^.

#### Identification of dLGN relay cells and TRN cells

We used both spike statistics and optogenetic tools to classify cells. Cells in TRN can be distinguished from relay cells by distinct burst inter-spike interval patterns^22,72–74^ (Figure S1B) and extracellular waveforms^47,50^. Photostimulation in PV-Ai32 mice activates cells in TRN but not in dLGN. Although the ventral division of LGN (vLGN) is also PV positive and lies near the posterior extreme of the visual sector of TRN^75^; cells there do not express T-type calcium channel and lack stereotypical bursts. Photostimulation in Gad2-Ai32 mice suppresses rather than activates relay cells and excites local interneurons.

#### Optogenetic stimulus

Optogenetic stimuli (450nm, ∼1mW at the probe tip) were generated using an Optogenetics-LED-Dual box (Prizmatix, Ltd.) and were delivered to the tip of the probe through lambda-B fiber (100-core, 1.2mm taper-tip) which was embedded to the electrode (Cambridge NeuroTech). LED pulses (25 ms at 5 Hz, 50 ms at 2 Hz, 100 ms at 1 Hz, or 250 ms at 1 Hz) were delivered within a 1s window, with pulse sequences generated using a Power1401 interface (Cambridge Electronic Design, Ltd.).

#### Visual Stimulus

Visual stimuli were generated using a ViSaGe MKII Stimulus Generator (Cambridge Research Systems Ltd.) and displayed on gamma corrected monitors (Acer V196L, refresh rate at 70Hz, mean luminance 180cd/m^2^). The monitors were mounted on adjustable arms and positioned to capture receptive fields in the contralateral hemifield. Each monitor was placed so that its midline was 180mm from the eye^17^; azimuth and elevation were measured subsequently.

The stimulus set included sparse noise, Gaussian (dense) noise, and natural movies^17,59,76,77^. Sparse noise consisted of black and white squares of different sizes (5°, 10°, 20°, and 30°) on a gray background. The squares were presented in pseudorandom order, 16 to 20 times each, at 256 positions along a 16×16 grid covering 80°×80° visual space. For a high-contrast sparse noise, the same sequence was used against a white or black background. Gaussian noise consisted of 3°, 5° or 7° squares, presented as “checkerboards” on grid sizes of 28, 16 or 12, respectively; the intensity of each square was drawn from a Gaussian distribution (33% RMS contrast). Gaussian noise sequences typically consisted of 16384 frames and were repeated 2-4 times. Unless specified otherwise, the stimulus rate was 35Hz (i.e., each frame of stimulus lasts for ∼28.5ms). Naturalistic movies^77^ were displayed at 100% contrast and repeated for 50 trials.

### Quantification and Statistical Analysis

#### Spike Sorting and Unit Selection

We used Kilosort4^78^ to sort spikes from signals recorded with multichannel electrodes. Kilosort outputs were manually curated with <u>Phy</u>. Single clusters that included multiunit activity that could not be separated cleanly into individual spikes were excluded. Clusters were merged if associated receptive fields were visually identical and there were no refractory period violations.

The position of each neuron relative to the tip of probe was associated with the mean x-y positions (*x*_kz_ and *y*_kz_ for horizontal and vertical displacement, respectively) of spikes generated by Kilosort4. Only cells having receptive fields within the stimulus grid were used for this study.

#### Histological Registration

Animals were perfused with 3% paraformaldehyde at the end of each experiment; the brain was removed and placed in 30% sucrose phosphate buffer and cut into 80 micron coronal sections with a sliding microtome (AO Scientific Instruments, Model 806). Tissue sections were mounted with ProLong Glass Antifade Mountant with NucBlue (ThermoFisher catalog#P36985) and imaged with a standard fluorescent microscope (Zeiss) or confocal microscope (Leica MICA) or slide scanner (Zeiss). Micrographs were first processed by DeepSlice^79^, which outputs a best-fitted linear transformation in Allen Mouse Brain Common Coordinate (CCFv3)^80^. The linear transformation was manually curated with QuickNII^81^, followed by nonlinear transformations using VisuAlign (RRID:SCR_017978).

Probe tracks from postmortem histology were used to localize recording sites. Pixel coordinates of the tip and the shaft of probe tracks were measured with custom scripts using FIJI^82^. In-house Python script was used to convert pixel coordinates into CCFv3 coordinates following the Bowyer-Watson algorithm specified by <u>VisuAl</u>ig n. The unit vector of the probe was estimated using singular value decomposition to represent its orientation. The coordinates of the shank tip (*x*_tip_, *y*_tip_, *z*_tip_, in CCFv3 AP, ML and DV coordinates) and its unit vector (*x*_u_, *y*_u_, *z*_u_) were used for the following analysis.

#### Anatomical locations of sorted units

Neuronal locations relative to the probe tip were estimated using Kilosort output (*x*_kz_ and *y*_kz_). Given that the width of each shank was usually less than 50μm, location coordinates (E, *Y* and *Z* for CCFv3 AP, ML and DV coordinates) were extrapolated only using the relative depth (*y*_kz_) within the probe (Figure S1A):

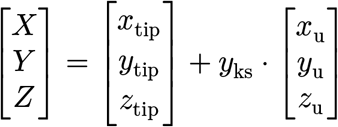

#### Visual receptive field estimation

Spatiotemporal receptive fields were estimated from responses to sparse noise or Gaussian noise by spike-triggered averaging (STA) of the stimulus ensemble^83^. Bright (On) and dark (Off) maps were generated for sparse noise by only using bright or dark squares, respectively; “difference” maps were made by subtracting dark from bright maps. STAs computed from Gaussian noise, in effect, yield difference maps^83^. We limited our analysis to windows as long as 300ms before each spike (i.e., ∼10 frames of stimuli displayed at 35Hz).

#### Spatial footprint of the receptive field

To quantify receptive fields without assumptions about their shape, we used a nonparametric footprint method, as described previously^22^. In essence, a Z-score for each pixel in the spatiotemporal STA was calculated by bootstrapping spike times; followed by a false discovery rate test to identify statistically significant pixels (Benjamini-Hochberg procedure, significance level < 0.01). The footprint of the receptive fields is thus defined as the time collapsed spatial map of the earliest significant pixels. Footprints of receptive fields from sparse noise were merged from separate estimations of the bright and dark maps; on-off pixels were defined as significant pixels in both bright and dark maps. Dominant pixels were selected as the polarity that elicited the stronger responses^22^. At population level, we found no significant differences between the raw estimation of sizes using footprint and standard 2D Gaussian fit, but measurement of individual cells differed (Figure S1C). Importantly, footprint analysis allowed us to project the receptive field straightforwardly onto the spherical representation of visuotopic space pixel-by-pixel.

#### Temporal course of receptive fields

We calculated the temporal STA (tSTA) by averaging the pixels within the spatial region delimited by the footprint for each stimulus frame, subtracting the baseline and normalizing to the absolute maximum, whether On or Off. The baseline was estimated as the mean response within a 300ms window **after** each spike. tSTAs computed from responses to sparse noise were calculated separately for bright and dark stimuli but normalized jointly.

Significant fluctuations in each tSTA were defined as the timepoints for which the Z-scored first-derivative exceeded ± 0.75 SD. For a biphasic tSTA with significant positive peak followed by a negative trough (or vice-versa), or a biphasic tSTA with two peaks of the same sign, the “triggering phase” was specified as the region bounded by spike time (t=0) and the zero-crossing or distinct inflection point; the “priming phase” was defined as the interval between the zero-crossing or distinct inflection point at an earlier time and a return to baseline. In cases, for which the tSTA had neither a detectable zero-crossing nor distinct inflection point, only the triggering phase was included in analysis. The triggering latency was defined as the first significant point of the triggering phase. The magnitude of the triggering or of the priming phase was defined as the integral of the entire phase; the ratio (priming-triggering ratio) between the magnitude of the priming phase and that of the triggering phase was used to assess the relative strength of the priming phase.

#### Classification of receptive-field types

We used responses to the smallest stimulus size that evoked a stable response to determine receptive field type. When the smallest effective stimulus size for sparse noise and Gaussian noise were the same, we conducted further measurements with sparse noise because it was possible to analyze bright and dark responses separately. Receptive field types were quantitatively grouped as On, Off, On-Off or Suppressed-by-Contrast (SbC) using the bright-dark polarity score (Ω) to tSTAs, as described previously^17,37^.

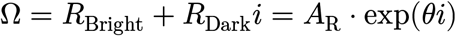

Where Ω is a complex number and *R*_Bright_ and *R*_Dark_ are values of bright and dark tSTA at the triggering latency of tSTAs. Ω can be rewritten into Euler’s formula, where the amplitude (*A*_R_) is the joint response level and angle (θ) points to the quadrant that represents a given receptive-field type: −180° < *θ* ≤ −90° indicates suppression by both bright and dark (Suppressed-by-Contrast, SbC); −90° < *θ* ≤ 0° indicates excitation by bright and suppression by dark (On); 0° < *θ* < 90° indicates excitation by both bright and dark (On-Off); 90° ≤ *θ* ≤ 180° indicates suppression by bright and excitation by dark (Off). Center-surround receptive fields (On-Center or Off-Center) were qualitatively classified from On and Off types afterwards as surrounds evoked by the smallest stimuli were often too weak to fit^16^.

#### Visuotopic projection of the spatial footprint

We estimated the visuotopic coordinates of each spatial footprint by projecting the flat monitor onto a unit sphere centered at the animal’s eye. The area of each grid in sparse noise and Gaussian noise is estimated as the solid angle on the unit sphere^84^. The Cartesian coordinates (**R**) of four corner pixels of the grid (***r****_a_*, ***r****_b_*, ***r****_c_*, ***r****_d_*) can be defined as:

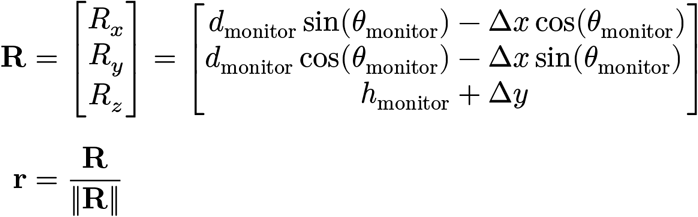

Where *R_x_* is the sagittal axis coordinates, *R_y_* is the horizontal axis coordinates and *R_z_* is the vertical axis coordinates. *θ*_monitor_ and ℎ_monitor_ are the azimuth and height of the monitor center relative to the eye, *d*_monitor_ is the distance between the midline of the monitor to the eye; Δ*x* and Δ*y* are the relative physical distance to the center of the monitor. The size of the square (S, in units of steradians) can be defined as the sum of two triangles:

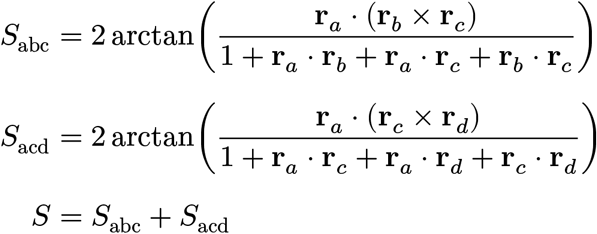

Receptive-field sizes were estimated as the sum of the corrected visual angles of the spherical projection and were converted to units of degrees squared. The center of the receptive field was estimated as the center of mass with smoothed weights using a 3×3 averaging filter. The visuotopic projection of the center (*θ* and *φ* for the azimuth and elevation) can be estimated as:

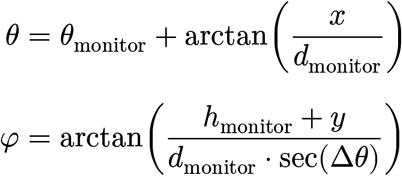

Where *x* and *y* are the physical location of the footprint center relative to the center of monitor; *d*_monitor_ is the distance between the midline of the monitor to the eye; *θ*_monitor_ and ℎ_monitor_ are the azimuth and height of the monitor.

#### Classification of receptive-field location

We grouped the centroids of receptive fields into binocular/monocular zones within a visuotopic reference frame. The putative optic axes of mouse eyes were chosen as ±64° in azimuth and 22° in elevation^4^. We assumed the retinal projection of the visual field spans 180°, illustrating the border of the left and right retinae (solid and dashed lines, respectively, Figure 3). All data were transformed with respect to the left retinal space. Thus, the spherical distance (*d*) from the receptive field center to the right retinal limits was used to assign the binocular versus monocular groups: 1) *d* < −5° as monocular regions; 2) *d* > 5° as binocular regions; 3) −5° ≤ *d* ≤ 5° as border regions and were excluded from statistical comparisons.

#### Burst and tonic spikes

Bursts were detected as described previously^21^. In brief, TRN bursts were identified as groups of 5 or more spikes within a 70 ms window that have interspike interval less than 30 ms and that follow a silent period of 70 ms or longer. Spike rate within a TRN burst typically accelerates and then decelerates (Figure S1B)^72^. For dLGN, bursts were identified as two or more spikes with maximum ISIs of 4 ms that follow a silent period of 100 ms or longer. Spikes within dLGN bursts decelerate over time. The first (cardinal) spike of each burst was used for burst analysis.

#### Statistics

All statistics and analysis were calculated using Julia^85^ and <u>“HypothesisTests.jl”</u> package. When two populations of non-parametric datasets were compared, we used the Mann-Whitney U test (`HypothesisTests.MannWhitneyUTest`); for non-parametric paired datasets, we used the Wilcoxon signed rank test (`HypothesisTests.SignedRankTest`). Unless specified, all statistical tests were two-tailed with sample size indicated in each figure and/or legend. Significance levels were indicated by asterisks: n.s., p ≥ 0.05; ∗, p < 0.05; ∗∗, p < 0.01; ∗∗∗, p < 0.001. Statistical variation for a given dataset is reported as mean ± SD.

## Data and Code Availability

The analyses of carnivore thalamus used previously published datasets^21,22^. The carnivore data for the figures in this paper are provided; requests for access to the original data should be directed to the corresponding author. The experimental data used in this paper will be available on Zenodo (<u>DOI: 10.5281/zenodo.21225776</u>). All code to generate results and figures is available upon request.

## Author contributions

YM, UMC and JAH initiated the research project and designed initial experimental strategies that were further developed by YM. YM and UMC conducted experiments in mouse TRN; YM, ASG, SA and RD conducted experiments in mouse dLGN; UMC and FTS developed initial methods to analyze bursts and YM developed all further analyses; UMC, RD and FTS contributed to analysis pipelines. YM and JAH wrote the manuscript and all other authors provided comments and approved the final text.

## Acknowledgments

The work was supported by NIH R01EY09395 to JAH and NIH R01EY037195 to FTS.

The authors declare no competing interests.

---

**Figure S1:**
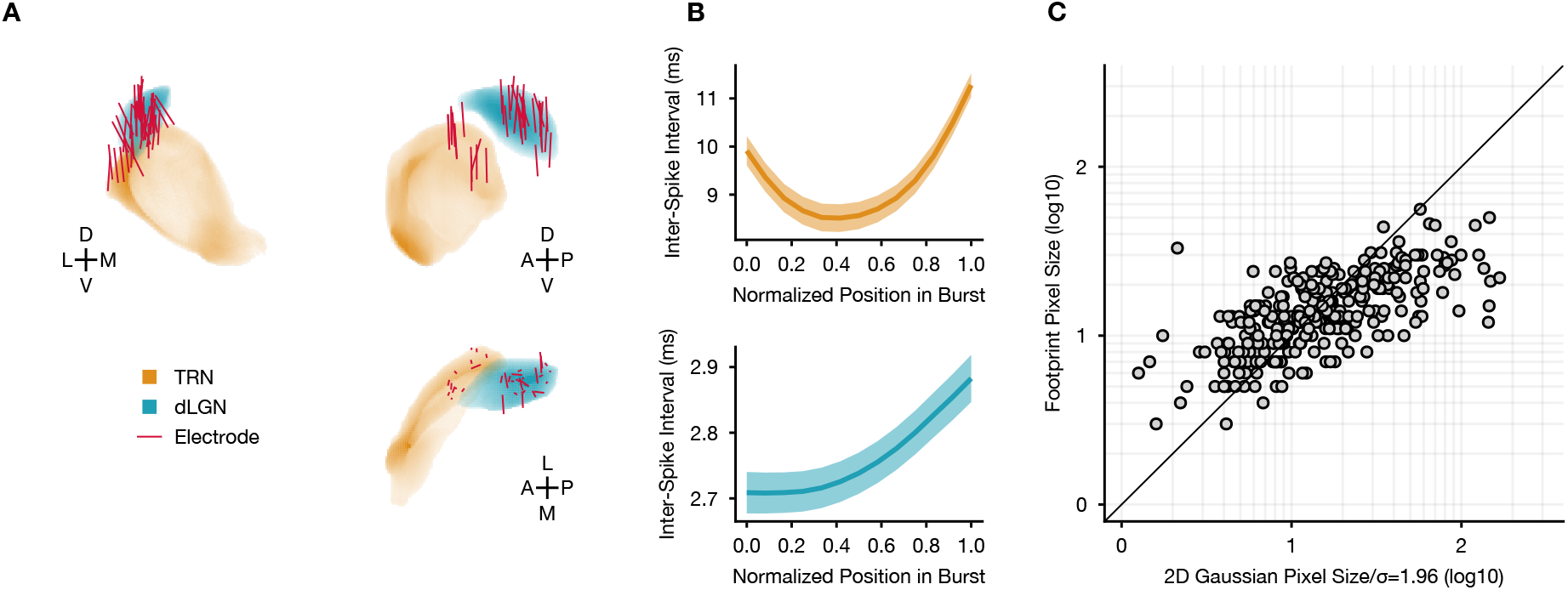
Identifying relay cells in dLGN and cells in TRN using *in vivo* recordings. (**A**) Three views of electrode tracks registered to Allen Mouse Brain Common Coordinate. D, dorsal; V, ventral; L, lateral; M, medial; A, anterior; P, posterior. **(B**) Averaged inter-spike interval (ISI) for bursts in TRN (Top) and dLGN (Bottom); the plot of ISIs is U-shaped for TRN but rises monotonically for dLGN. **(C**) Scatter plots for estimations of receptive-field sizes (**not** corrected for visuotopic projection) using 2D Gaussian fit (1.96σ) and footprint analysis; diagonal line shows unity slope.

**Figure S2:**
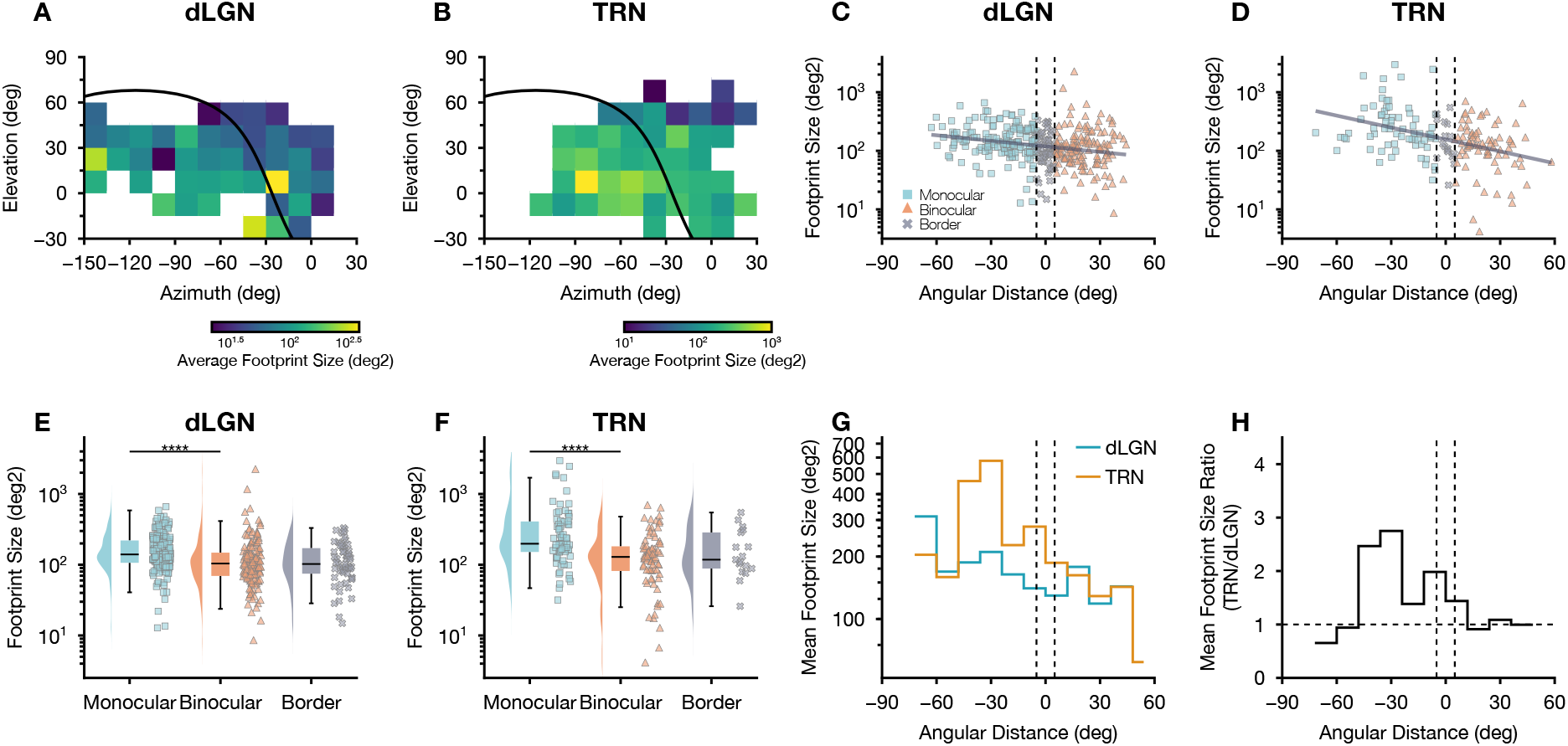
Non-uniform distribution of dominant footprint sizes in murine dLGN and TRN across visuotopic space. (**A, B**) Heatmap of averaged footprint sizes (using dominant pixels) across visuotopic space (bin size, 15°) for cells in dLGN (A) and TRN (B) from Figure 4. **(C, D**) Scatter plots show the relationship of footprint sizes (using dominant pixels) from dLGN (C) and TRN (D) plotted on a log scale against the angular distance from the border of the monocular visual field, with the linear regression for each population plotted as solid line. Vertical dashed lines indicate the border between binocular and monocular zones. (F-test; dLGN, p=4.523 × 10^−7^; TRN, p=2.149 × 10^−5^). **(E, F**) Raincloud plots show ranges of footprint sizes (using dominant pixels) in the monocular, binocular and border regions for cells in dLGN (E, Mann-Whitney U test, p=1.763 × 10^−7^) and TRN (F, Mann-Whitney U test, p=8.141 × 10^−6^) **(G**) Histograms plot distributions of averaged footprint sizes (using dominant pixels) of cells in dLGN and TRN along angular distance from the border of monocular visual field; bin size, 12°. **(H**) Histograms of ratios between the averaged footprint sizes (using dominant pixels) of dLGN cells and TRN cells at each angular distance. The horizontal dotted line indicates ratio=1.

**Figure S3:**
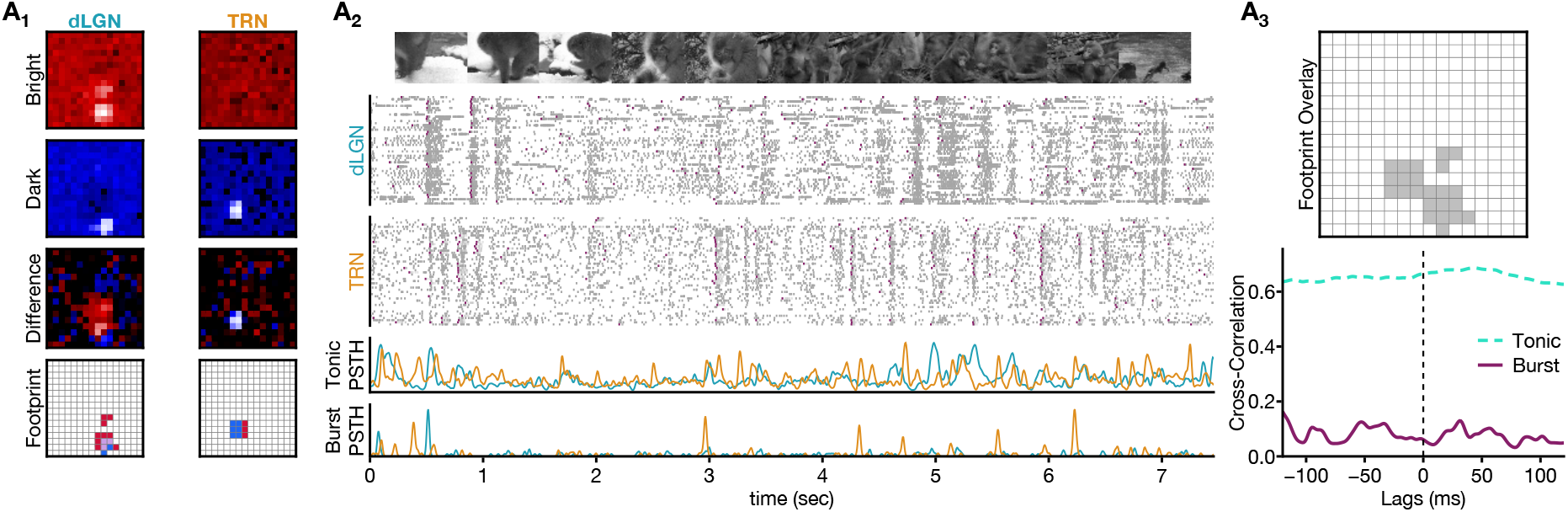
Bursts in TRN did not precede bursts in relay cells with displaced receptive fields. Conventions as in Figure 7, but for example responses from the dLGN On-Off cell in Figure 7B and a TRN On-Off cell with displaced receptive fields (gray, non-overlapping pixels).

## References

1. Pinault, D., and Deschênes, M. (1998). Projection and innervation patterns of individual thalamic reticular axons in the thalamus of the adult rat: A three-dimensional, graphic, and morphometric analysis. J. Comp. Neurol. 391, 180–203. 10.1002/(sici)1096-9861(19980209)391:2<180::aid-cne3>3.0.co;2-z.

2. Pinault, D. (2004). The thalamic reticular nucleus: Structure, function and concept. Brain Res. Rev. 46, 1–31. 10.1016/j.brainresrev.2004.04.008.

3. Baden, T., Euler, T., and Berens, P. (2020). Understanding the retinal basis of vision across species. Nat. Rev. Neurosci. 21, 5–20. 10.1038/s41583-019-0242-1.

4. Sterratt, D.C., Lyngholm, D., Willshaw, D.J., and Thompson, I.D. (2013). Standard anatomical and visual space for the mouse retina: Computational reconstruction and transformation of flattened retinae with the Retistruct package. PLOS Comput. Biol. 9, e1002921. 10.1371/journal.pcbi.1002921.

5. Heukamp, A.S., Warwick, R.A., and Rivlin-Etzion, M. (2020). Topographic variations in retinal encoding of visual space. Annu. Rev. Vision Sci. 6, 237–259. 10.1146/annurev-vision-121219-081831.

6. Budoff, S.A., and Poleg-Polsky, A. (2025). A complete spatial map of mouse retinal ganglion cells reveals density and gene expression specializations. Proc. Natl. Acad. Sci. 122, e2515449122. 10.1073/pnas.2515449122.

7. Gallego-Ortega, A., Norte-Muñoz, M., Di Pierdomenico, J., Avilés-Trigueros, M., De La Villa, P., Valiente-Soriano, F.J., and Vidal-Sanz, M. (2022). Alpha retinal ganglion cells in pigmented mice retina: Number and distribution. Front. Neuroanat. 16, 1054849. 10.3389/fnana.2022.1054849.

8. Bleckert, A., Schwartz, G., Turner, M., Rieke, F., and Wong, R. (2014). Visual space is represented by nonmatching topographies of distinct mouse retinal ganglion cell types. Curr. Biol. 24, 310–315. 10.1016/j.cub.2013.12.020.

9. Sanes, J.R., and Masland, R.H. (2015). The types of retinal ganglion cells: Current status and implications for neuronal classification. Annu. Rev. Neurosci. 38, 221–246. 10.1146/annurev-neuro-071714-034120.

10. Kerschensteiner, D. (2022). Feature detection by retinal ganglion cells. Annu. Rev. Vision Sci. 8, 135–169. 10.1146/annurev-vision-100419-112009.

11. Baden, T., Berens, P., Franke, K., Román Rosón, M., Bethge, M., and Euler, T. (2016). The functional diversity of retinal ganglion cells in the mouse. Nature 529, 345–350. 10.1038/nature16468.

12. Gupta, D., Młynarski, W., Sumser, A., Symonova, O., Svatoň, J., and Joesch, M. (2023). Panoramic visual statistics shape retina-wide organization of receptive fields. Nat. Neurosci. 26, 606–614. 10.1038/s41593-023-01280-0.

13. Skyberg, R.J., and Niell, C.M. (2024). Natural visual behavior and active sensing in the mouse. Curr. Opin. Neurobiol. 86, 102882. 10.1016/j.conb.2024.102882.

14. Johnson, K.P., Fitzpatrick, M.J., Zhao, L., Wang, B., McCracken, S., Williams, P.R., and Kerschensteiner, D. (2021). Cell-type-specific binocular vision guides predation in mice. Neuron 109, 1527–1539.e4. 10.1016/j.neuron.2021.03.010.

15. Kerschensteiner, D., and Guido, W. (2017). Organization of the dorsal lateral geniculate nucleus in the mouse. Visual Neurosci. 34, E008. 10.1017/S0952523817000062.

16. Piscopo, D.M., Huberman, A.D., and Niell, C.M. (2013). Diverse visual features encoded in mouse lateral geniculate nucleus. J. Neurosci. 33, 4642–4656. 10.1523/JNEUROSCI.5187-12.2013.

17. Suresh, V., Ciftcioglu, U.M., Wang, X., Lala, B.M., Ding, K.R., Smith, W.A., Sommer, F.T., and Hirsch, J.A. (2016). Synaptic contributions to receptive field structure and response properties in the rodent lateral geniculate nucleus of the thalamus. J. Neurosci. 36, 10949–10963. 10.1523/JNEUROSCI.1045-16.2016.

18. Usrey, W.M., Reppas, J.B., and Reid, R.C. (1999). Specificity and strength of retinogeniculate connections. J. Neurophysiol. 82, 3527–3540. 10.1152/jn.1999.82.6.3527.

19. Xue, J., Carney, T., Ramoa, A., and Freeman, R. (1988). Binocular interaction in the perigeniculate nucleus of the cat. Exp. Brain Res. 69. 10.1007/BF00247304.

20. Usrey, W.M., and Alitto, H.J. (2015). Visual functions of the thalamus. Annu. Rev. Vision Sci. 1, 351–371. 10.1146/annurev-vision-082114-035920.

21. Vaingankar, V., Sanchez Soto, C., Wang, X., Sommer, F.T., and Hirsch, J.A. (2012). Neurons in the thalamic reticular nucleus are selective for diverse and complex visual features. Front. Integr. Neurosci. 6. 10.3389/fnint.2012.00118.

22. Soto-Sánchez, C., Wang, X., Vaingankar, V., Sommer, F.T., and Hirsch, J.A. (2017). Spatial scale of receptive fields in the visual sector of the cat thalamic reticular nucleus. Nat. Commun. 8, 800. 10.1038/s41467-017-00762-7.

23. Jahnsen, H., and Llinás, R. (1984). Electrophysiological properties of Guinea-pig thalamic neurones: An in vitro study. J. Physiol. 349, 205–226. 10.1113/jphysiol.1984.sp015153.

24. Huguenard, J., and Prince, D. (1992). A novel T-type current underlies prolonged Ca(2+)-dependent burst firing in GABAergic neurons of rat thalamic reticular nucleus. J. Neurosci. 12, 3804–3817. 10.1523/JNEUROSCI.12-10-03804.1992.

25. Lesica, N.A., Weng, C., Jin, J., Yeh, C.-I., Alonso, J.-M., and Stanley, G.B. (2006). Dynamic encoding of natural luminance sequences by LGN bursts. PLOS Biol. 4, e209. 10.1371/journal.pbio.0040203.

26. Denning, K.S., and Reinagel, P. (2005). Visual control of burst priming in the anesthetized lateral geniculate nucleus. J. Neurosci. 25, 3531–3538. 10.1523/JNEUROSCI.4417-04.2005.

27. Wang, X., Wei, Y., Vaingankar, V., Wang, Q., Koepsell, K., Sommer, F.T., and Hirsch, J.A. (2007). Feedforward excitation and inhibition evoke dual modes of firing in the cat’s visual thalamus during naturalistic viewing. Neuron 55, 465–478. 10.1016/j.neuron.2007.06.033.

28. Alitto, H.J., Weyand, T.G., and Usrey, W.M. (2005). Distinct properties of stimulus-evoked bursts in the lateral geniculate nucleus. J. Neurosci. 25, 514–523. 10.1523/JNEUROSCI.3369-04.2005.

29. Crick, F. (1984). Function of the thalamic reticular complex: The searchlight hypothesis. Proc. Natl. Acad. Sci. 81, 4586–4590. 10.1073/pnas.81.14.4586.

30. McAlonan, K., Cavanaugh, J., and Wurtz, R.H. (2008). Guarding the gateway to cortex with attention in visual thalamus. Nature 456, 391–394. 10.1038/nature07382.

31. Halassa, M.M., and Kastner, S. (2017). Thalamic functions in distributed cognitive control. Nat. Neurosci. 20, 1669–1679. 10.1038/s41593-017-0020-1.

32. Vaughn, M.J., Laswick, Z., Wang, H., and Haas, J.S. (2024). Functionally distinct circuits are linked by heterocellular electrical synapses in the thalamic reticular nucleus. eNeuro 11, ENEURO.0269–23.2023. 10.1523/ENEURO.0263-23.2023.

33. Cho, F.S., Huguenard, J.R., and Paz, J.T. (2025). A dual inhibitory network in the thalamic reticular nucleus delineated by pallidal and intra-reticular inhibition. Preprint at bioRxiv, 10.1101/2025.10.05.680571.

34. Fitzgibbon, T., Szmajda, B.A., and Martin, P.R. (2007). First order connections of the visual sector of the thalamic reticular nucleus in marmoset monkeys (Callithrix jacchus). Visual Neurosci. 24, 857–874. 10.1017/S0952523807070770.

35. Wang, S., Bickford, M.E., Horn, S.C. van, Erisir, A., Godwin, D.W., and Sherman, S.M. (2001). Synaptic targets of thalamic reticular nucleus terminals in the visual thalamus of the cat. J. Comp. Neurol. 440, 321–341. 10.1002/cne.1383.

36. Boshra, R., Harris, M., Dougherty, K., Berg, M., Morea, B.M., Alitto, H.J., Rodriguez, M.C., Usrey, W.M., and Kastner, S. (2025). Causal role for pulvinar burst firing in thalamo-cortical attention control. Preprint at bioRxiv, 10.1101/2025.09.16.676591.

37. Gorin, A.S., Miao, Y., Ahn, S., Suresh, V., Su, Y., Ciftcioglu, U.M., Sommer, F.T., and Hirsch, J.A. (2023). Local interneurons in the murine visual thalamus have diverse receptive fields and can provide feature selective inhibition to relay cells. Preprint at bioRxiv, 10.1101/2023.08.10.549394.

38. Grubb, M.S., and Thompson, I.D. (2003). Quantitative characterization of visual response properties in the mouse dorsal lateral geniculate nucleus. J. Neurophysiol. 90, 3594–3607. 10.1152/jn.00633.2003.

39. McAlonan, K. (2006). Attentional modulation of thalamic reticular neurons. J. Neurosci. 26, 4444–4450. 10.1523/JNEUROSCI.5602-05.2006.

40. Goetz, J., Jessen, Z.F., Jacobi, A., Mani, A., Cooler, S., Greer, D., Kadri, S., Segal, J., Shekhar, K., Sanes, J.R., et al. (2022). Unified classification of mouse retinal ganglion cells using function, morphology, and gene expression. Cell Rep. 40, 111040. 10.1016/j.celrep.2022.111040.

41. Alitto, H., Rathbun, D.L., Vandeleest, J.J., Alexander, P.C., and Usrey, W.M. (2019). The augmentation of retinogeniculate communication during thalamic burst mode. J. Neurosci. 39, 5697–5710. 10.1523/JNEUROSCI.2320-18.2019.

42. Lesica, N.A., and Stanley, G.B. (2004). Encoding of natural scene movies by tonic and burst spikes in the lateral geniculate nucleus. J. Neurosci. 24, 10731–10740. 10.1523/JNEUROSCI.3059-04.2004.

43. Bragg, E.M., Fairless, E.A., Liu, S., and Briggs, F. (2017). Morphology of visual sector thalamic reticular neurons in the macaque monkey suggests retinotopically specialized, parallel stream-mixed input to the lateral geniculate nucleus. J. Comp. Neurol. 525, 1273–1290. 10.1002/cne.24134.

44. Crabtree, J.W. (1999). Intrathalamic sensory connections mediated by the thalamic reticular nucleus. Cell. Mol. Life Sci. 56, 683–700. 10.1007/s000180050462.

45. Baldauf, Z. (2010). Dual chemoarchitectonic lamination of the visual sector of the thalamic reticular nucleus. Neuroscience 165, 801–818. 10.1016/j.neuroscience.2003.11.010.

46. Crabtree, J.W., and Killackey, H.P. (1989). The topographic organization and axis of projection within the visual sector of the rabbit’s thalamic reticular nucleus. Eur. J. Neurosci. 1, 94–109. 10.1111/j.1460-9568.1989.tb00777.x.

47. Li, Y., Lopez-Huerta, V.G., Adiconis, X., Levandowski, K., Choi, S., Simmons, S.K., Arias-Garcia, M.A., Guo, B., Yao, A.Y., Blosser, T.R., et al. (2020). Distinct subnetworks of the thalamic reticular nucleus. Nature 583, 819–824. 10.1038/s41586-020-2504-5.

48. Martinez-Garcia, R.I., Voelcker, B., Zaltsman, J.B., Patrick, S.L., Stevens, T.R., Connors, B.W., and Cruikshank, S.J. (2020). Two dynamically distinct circuits drive inhibition in the sensory thalamus. Nature 583. 10.1038/s41586-020-2512-5.

49. Pinault, D., and Deschênes, M. (1998). Anatomical evidence for a mechanism of lateral inhibition in the rat thalamus. Eur. J. Neurosci. 10, 3462–3469. 10.1046/j.1460-9568.1998.00362.x.

50. Hartley, N.D., Krol, A., Choi, S., Rome, N., Levandowski, K., Pasqualoni, S., Jones, C., Tian, J., Lee, S., Lee, H., et al. (2024). Distinct structural and functional connectivity of genetically segregated thalamoreticular subnetworks. Cell Rep. 43, 115037. 10.1016/j.celrep.2024.115037.

51. Clemente-Perez, A., Makinson, S.R., Higashikubo, B., Brovarney, S., Cho, F.S., Urry, A., Holden, S.S., Wimer, M., Dávid, C., Fenno, L.E., et al. (2017). Distinct thalamic reticular cell types differentially modulate normal and pathological cortical rhythms. Cell Rep. 19, 2130–2142. 10.1016/j.celrep.2017.05.044.

52. Pinault, D., Bourassa, J., and Deschênes, M. (1995). Thalamic reticular input to the rat visual thalamus: A single fiber study using biocytin as an anterograde tracer. Brain Res. 670, 147–152. 10.1016/0006-8993(94)01303-Y.

53. Connolly, M., and Van Essen, D. (1984). The representation of the visual field in parvicellular and magnocellular layers of the lateral geniculate nucleus in the macaque monkey. J. Comp. Neurol. 226, 544–564. 10.1002/cne.902260408.

54. Sanderson, K.J. (1971). The projection of the visual field to the lateral geniculate and medial interlaminar nuclei in the cat. J. Comp. Neurol. 143, 101–117. 10.1002/cne.901430107.

55. Hoy, J.L., Yavorska, I., Wehr, M., and Niell, C.M. (2016). Vision drives accurate approach behavior during prey capture in laboratory mice. Curr. Biol. 26, 3046–3052. 10.1016/j.cub.2016.03.003.

56. Denman, D.J., Siegle, J.H., Koch, C., Reid, R.C., and Blanche, T.J. (2017). Spatial organization of chromatic pathways in the mouse dorsal lateral geniculate nucleus. J. Neurosci. 37, 1102–1116. 10.1523/JNEUROSCI.1742-16.2016.

57. Van Beest, E.H., Mukherjee, S., Kirchberger, L., Schnabel, U.H., Van Der Togt, C., Teeuwen, R.R.M., Barsegyan, A., Meyer, A.F., Poort, J., Roelfsema, P.R., et al. (2021). Mouse visual cortex contains a region of enhanced spatial resolution. Nat. Commun. 12, 4029. 10.1038/s41467-021-24311-5.

58. Born, G., Schneider-Soupiadis, F.A., Erisken, S., Vaiceliunaite, A., Lao, C.L., Mobarhan, M.H., Spacek, M.A., Einevoll, G.T., and Busse, L. (2021). Corticothalamic feedback sculpts visual spatial integration in mouse thalamus. Nat. Neurosci. 24, 1711–1720. 10.1038/s41593-021-00943-0.

59. Wang, X., Vaingankar, V., Soto Sanchez, C., Sommer, F.T., and Hirsch, J.A. (2011). Thalamic interneurons and relay cells use complementary synaptic mechanisms for visual processing. Nat. Neurosci. 14, 224–231. 10.1038/nn.2707.

60. Bullier, J.H., and Norton, T.T. (1977). Receptive-field properties of X-, Y– and intermediate cells in the cat lateral geniculate nucleus. Brain Res. 121, 151–156. 10.1016/0006-8993(77)90444-9.

61. Uhlrich, D.J., Cucchiaro, J.B., Humphrey, A.L., and Sherman, S.M. (1991). Morphology and axonal projection patterns of individual neurons in the cat perigeniculate nucleus. J. Neurophysiol. 65, 1528–1541. 10.1152/jn.1991.65.6.1528.

62. Conley, M., and Diamond, I.T. (1990). Organization of the visual sector of the thalamic reticular nucleus in Galago: Evidence that the dorsal lateral geniculate and pulvinar nuclei occupy separate parallel tiers. Eur. J. Neurosci. 2, 211–226. 10.1111/j.1460-9568.1990.tb00414.x.

63. Destexhe, A., and Sejnowski, T.J. (2002). The initiation of bursts in thalamic neurons and the cortical control of thalamic sensitivity. Philos. Trans. R. Soc. Lond., B: Biol. Sci. 357, 1649–1657. 10.1098/rstb.2002.1154.

64. O’Malley, J.J., Seibt, F., Chin, J., and Beierlein, M. (2020). TRPM4 conductances in thalamic reticular nucleus neurons generate persistent firing during slow oscillations. J. Neurosci. 40, 4813–4823. 10.1523/JNEUROSCI.0324-20.2020.

65. Crabtree, J.W. (2018). Functional diversity of thalamic reticular subnetworks. Front. Syst. Neurosci. 12, 41. 10.3389/fnsys.2018.00041.

66. Lee, S.-C., Patrick, S.L., Richardson, K.A., and Connors, B.W. (2014). Two functionally distinct networks of gap junction-coupled inhibitory neurons in the thalamic reticular nucleus. J. Neurosci. 34, 13170–13182. 10.1523/JNEUROSCI.0562-14.2014.

67. Hoseini, M.S., Higashikubo, B., Cho, F.S., Chang, A.H., Clemente-Perez, A., Lew, I., Ciesielska, A., Stryker, M.P., and Paz, J.T. (2021). Gamma rhythms and visual information in mouse V1 specifically modulated by somatostatin+ neurons in reticular thalamus. eLife 10, e61437. 10.7554/eLife.61437.

68. Whilden, C.M., Chevée, M., An, S.Y., and Brown, S.P. (2021). The synaptic inputs and thalamic projections of two classes of layer 6 corticothalamic neurons in primary somatosensory cortex of the mouse. J. Comp. Neurol. 529, 3751–3771. 10.1002/cne.25163.

69. Carroll, B.J., Sampathkumar, V., Kasthuri, N., and Sherman, S.M. (2022). Layer 5 of cortex innervates the thalamic reticular nucleus in mice. Proc. Natl. Acad. Sci. 119, e2205209119. 10.1073/pnas.2205209119.

70. Usrey, W., and Fitzpatrick, D. (1996). Specificity in the axonal connections of layer VI neurons in tree shrew striate cortex: Evidence for distinct granular and supragranular systems. J. Neurosci. 16, 1203–1218. 10.1523/JNEUROSCI.16-03-01203.1996.

71. Kirchgessner, M.A., Franklin, A.D., and Callaway, E.M. (2020). Context-dependent and dynamic functional influence of corticothalamic pathways to first– and higher-order visual thalamus. Proc. Natl. Acad. Sci. 117, 13066–13077. 10.1073/pnas.2002080117.

72. Domich, L., Oakson, G., and Steriade, M. (1986). Thalamic burst patterns in the naturally sleeping cat: A comparison between cortically projecting and reticularis neurones. J. Physiol. 379, 429–449. 10.1113/jphysiol.1986.sp016262.

73. Kimura, A., Yokoi, I., Imbe, H., Donishi, T., and Kaneoke, Y. (2012). Distinctions in burst spiking between thalamic reticular nucleus cells projecting to the dorsal lateral geniculate and lateral posterior nuclei in the anesthetized rat. Neuroscience 226, 208–226. 10.1016/j.neuroscience.2012.03.016.

74. Fernandez, L.M., Vantomme, G., Osorio-Forero, A., Cardis, R., Béard, E., and Lüthi, A. (2018). Thalamic reticular control of local sleep in mouse sensory cortex. eLife 7, e39111. 10.7554/eLife.33111.

75. Sabbagh, U., Govindaiah, G., Somaiya, R.D., Ha, R.V., Wei, J.C., Guido, W., and Fox, M.A. (2021). Diverse GABAergic neurons organize into subtype-specific sublaminae in the ventral lateral geniculate nucleus. J. Neurochem. 159, 479–497. 10.1111/jnc.15101.

76. Wang, Q., and Burkhalter, A. (2007). Area map of mouse visual cortex. J. Comp. Neurol. 502, 339–357. 10.1002/cne.21286.

77. Ciftcioglu, U.M., Suresh, V., Ding, K.R., Sommer, F.T., and Hirsch, J.A. (2020). Visual information processing in the ventral division of the mouse lateral geniculate nucleus of the thalamus. J. Neurosci. 40, 5019–5032. 10.1523/JNEUROSCI.2602-19.2020.

78. Pachitariu, M., Sridhar, S., Pennington, J., and Stringer, C. (2024). Spike sorting with Kilosort4. Nat. Methods 21, 914–921. 10.1038/s41592-024-02232-7.

79. Carey, H., Pegios, M., Martin, L., Saleeba, C., Turner, A.J., Everett, N.A., Bjerke, I.E., Puchades, M.A., Bjaalie, J.G., and McMullan, S. (2023). DeepSlice: Rapid fully automatic registration of mouse brain imaging to a volumetric atlas. Nat. Commun. 14, 5884. 10.1038/s41467-023-41645-4.

80. Wang, Q., Ding, S.-L., Li, Y., Royall, J., Feng, D., Lesnar, P., Graddis, N., Naeemi, M., Facer, B., Ho, A., et al. (2020). The Allen Mouse Brain Common Coordinate Framework: A 3D reference atlas. Cell 181, 936–953.e20. 10.1016/j.cell.2020.04.007.

81. Puchades, M.A., Csucs, G., Ledergerber, D., Leergaard, T.B., and Bjaalie, J.G. (2019). Spatial registration of serial microscopic brain images to three-dimensional reference atlases with the QuickNII tool. PLOS One 14, e0216796. 10.1371/journal.pone.0216736.

82. Schindelin, J., Arganda-Carreras, I., Frise, E., Kaynig, V., Longair, M., Pietzsch, T., Preibisch, S., Rueden, C., Saalfeld, S., Schmid, B., et al. (2012). Fiji: An open-source platform for biological-image analysis. Nat. Methods 9, 676–682. 10.1038/nmeth.2013.

83. Schwartz, O., Pillow, J.W., Rust, N.C., and Simoncelli, E.P. (2006). Spike-triggered neural characterization. J. Vis. 6, 484–507. 10.1167/6.4.13.

84. Van Oosterom, A., and Strackee, J. (1983). The solid angle of a plane triangle. IEEE Trans. Bio-Med. Eng. BME*-*30, 125–126. 10.1109/TBME.1983.325207.

85. Bezanson, J., Edelman, A., Karpinski, S., and Shah, V.B. (2017). Julia: A fresh approach to numerical computing. SIAM Rev. 59, 65–98. 10.1137/141000671.

